# Morphological Subprofile Analysis for Bioactivity Annotation of Small Molecules

**DOI:** 10.1101/2022.08.15.503944

**Authors:** Axel Pahl, Beate Schölermann, Marion Rusch, Mark Dow, Christian Hedberg, Adam Nelson, Sonja Sievers, Herbert Waldmann, Slava Ziegler

## Abstract

Fast prediction of mode of action for bioactive compounds would immensely foster bioactivity annotation in compound collections and may early on reveal off-targets in chemical biology research and drug discovery. A variety of target-based assays is available for addressing the modulation of druggable proteins. However, they cannot precisely predict how a compound would influence cellular processes due to polypharmacology. Furthermore, non-protein targets are often not considered. Morphological profiling, e.g., using the Cell Painting assay that monitors hundreds of morphological features upon compound perturbation and staining of cellular components, offers a fast, unbiased assessment of compound activity on various targets and cellular processes in one single experiment. However, due to incomplete bioactivity annotation and unknown activities of reference (landmark) compounds, prediction of bioactivity is not straightforward. Here we introduce the concept of subprofile analysis to map the mode of action for both reference and unexplored compounds. We defined mode-of-action clusters for a group of reference compounds and extracted cluster subprofiles that contain only a subset of morphological features (i.e., subprofiles) to represent a consensus profile. Subprofile analysis allows for assignment of compounds to, currently, ten different targets or modes of action in one single assay and bypasses the need of exploring all biosimilar reference compounds for the generation of target hypothesis. This approach will enable rapid bioactivity annotation of compound collections, particularly of uncharacterized small molecules, and will be extended to further bioactivity clusters in future. The data is public accessible via https://github.com/mpimp-comas/2022_pahl_ziegler_subprofiles and the web app tool http://cpcse.pythonanywhere.com/.

## Introduction

The design, synthesis and biological investigation of compound collections is at the heart of chemical biology research and drug discovery. Target- and cell-based assays are frequently employed to detect modulators of disease-relevant targets or processes. However, these assays are usually biased towards the kind of the sought bioactivity. Profiling approaches, which monitor hundreds of parameters like gene or protein expression or morphological features, provide a broader view on biological states in cells and organisms (Ziegler et al., 2021). Profiling of small molecules in cells provides an unbiased snapshot of perturbed processes and may uncover novel and off-targets. Morphological profiling, e.g., using the Cell Painting assay (CPA), monitors the change in morphological features, has higher throughput than gene and protein expression profiling and has been used to detect bioactivity in compound collections (Ziegler et al., 2021). The CPA employs six different dyes for detection of cell organelles and components (DNA, RNA, mitochondria, Golgi, plasma membrane, endoplasmic reticulum, actin cytoskeleton) (Bray et al., 2016, Gustafsdottir et al., 2013). Generation of CPA profiles for reference compounds is crucial and a prerequisite for the analysis of bioactivity of uncharacterized small molecules. Ideally, reference compounds sharing the same target shall give rise to similar CPA profiles and profile similarity can then be used to predict a target for a new compound. However, reference compounds often lack complete annotation, may display polypharmacology and address unknown targets which significantly hampers the generation of target hypotheses based solely on profile similarity (Akbarzadeh et al., 2021, Schneidewind et al., 2020, Schneidewind et al., 2021).

Herein we introduce the concept of subprofile analysis to define cluster subprofiles for bioactivity clusters that are detectable using the Cell Painting assay. These cluster subprofiles consist only of features that are common to reference compounds within one bioactivity cluster. Similarity to the cluster subprofiles is then sufficient to reliably predict a mode of action (MoA) related to biological targets such as AKT/PI3K/MTOR, Aurora kinases, BET, DNA synthesis, HDAC, HSP90 and tubulin or processes like lysosomotropism/cholesterol homeostasis regulation, protein synthesis and uncoupling of the mitochondrial proton gradient. Non-cluster features, i.e., features from the full profile that do not belong to the cluster subprofile, can in addition be employed to map activity that is not related to the cluster features or to group compounds within one MoA cluster. The subprofile approach allows for easy and rapid annotation of bioactivity of unexplored compounds using the Cell Painting assay without prior knowledge of the top biosimilar reference compounds and can detect polypharmacology.

## Results

For hypothesis generation in the biological analysis of our in-house compound collection, we explored a set of 4,251 reference compounds in a wide range of concentrations (10,551 measurements) using the CPA. Therefore, U2OS cells were treated with the compounds for 20 h prior to staining of cell compartments and components such as DNA, RNA, actin, plasma membrane, Golgi, mitochondria and endoplasmic reticulum (Bray et al., 2016, Gustafsdottir et al., 2013). Morphological profiles consist of Z scores of 579 features, which represent the difference for each feature relative to the DMSO control. The percentage of altered features defined as the induction (in %) is used as a measure of bioactivity and compounds are considered active for induction ≥ 5%. Profile similarity (termed biosimilarity, in %) is used for profile comparison and profiles are similar to each other if biosimilarity is higher than 75 %. Of the 4,162 references tested in a concentration range between 2 and 10 µM, 1,472 (35%) displayed activity with an induction ≥ 5 %. The final data set used is composed of 1,883 different reference compounds and three internal compounds (in total 3,560 measurements at different concentrations) that were non-toxic (cell count > 50 % compared to the DMSO control) and showed an induction between 5 and 85 %. Analysis of this data set revealed only in few cases profile similarity for compounds with similar target annotation, e.g., for inhibitors of Aurora kinase, BET, HDAC, HSP90 and protein synthesis. Instead, for a given reference compound, we detected diverse annotated activities for the most highly biosimilar compounds. Whereas biosimilarity of MTOR inhibitors to PI3K and AKT is most likely attributed to modulating the same pathway (AKT/PI3K/MTOR), we recently detected biosimilarity of the iron chelator deferoxamine to nucleoside analogues, antifolates and CDK inhibitors (Schneidewind et al., 2020). The biosimilarity of these compounds is not attributed to impairment of the same target but rather the same process, namely DNA synthesis and, thus, the cell cycle. Furthermore, we identified several reference compounds that share similar CPA profiles with Nocodazole, a well-studied tubulin inhibitor, although for most of these compounds the bioactivity annotation was different from tubulin targeting (Akbarzadeh et al., 2021). A literature search revealed impairment of microtubules for several compounds and we experimentally validated tubulin targeting for reference compounds that had not been previously linked to tubulin. In addition, 12 % of the reference set (27 % of the active references) belong to a large cluster with deregulation of cholesterol biosynthesis as a common denominator (Schneidewind et al., 2021). This cluster contains compounds with very diverse targets and the activity of approx. 75 % of the compounds is most likely due to their physicochemical properties that lead to accumulation of compounds in lysosomes due to protonation, thereby raising lysosomal pH and affecting the activity of lysosomes, lysosomal enzymes and lysosome-dependent processes such as autophagy. We also detected a cluster of biosimilar compounds related to the profile of the uncoupling reagent FCCP (see Figure S1). FCCP is biosimilar to Tyrphostin AG 879 and Tyrphostin A9 that are known to be kinase inhibitors. However, tyrphostins also disturb the mitochondrial proton gradient (Soltoff, 2004, Childress et al., 2018). Moreover, the antimicrobial agent triclosan and the IKK kinase inhibitor IMD0354 were biosimilar to FCCP and uncoupling activity for these compounds has been reported (Weatherly et al., 2016, Lee et al., 2013) although this information rarely is included in the compound annotation. Thus, Cell Painting detects uncoupling of the mitochondrial proton gradient as the common mode of action for these small molecules, as observed also by different morphological profiling approaches (Ziegler et al., 2021). Consequently, for the CPA profile of an uncharacterized small molecule and even a reference compound, a simple inspection of the top biosimilar reference compounds often would not lead to generation of target hypothesis as frequently compounds are biosimilar to several apparently not related reference compounds. One possibility is to map the location of the profile in a lower dimension plot (e.g., PCA, tSNE or UMAP), however, bioactivity clusters have to be defined beforehand and, more importantly, polypharmacological compounds may fail to localize in a separate cluster as recently shown for the BET inhibitor CF53 (Akbarzadeh et al., 2021).

To enhance the correct assignment to bioactivity (MoA) clusters, we set out to determine the characteristic profile of each cluster that potentially could be employed to assess biosimilarity to a predefined cluster rather than to single references. As compounds change only a subset of the 579 morphological features, we envisioned that we can extract these altered features for a given bioactivity cluster to yield a cluster ‘subprofile’. If such subprofiles represented the MoA of the clusters, they could successfully be used to simultaneously map similarity of compounds to all clusters defined thus far. To generate such subprofiles, first the dominating features for a set of biosimilar compounds are extracted and then a representative consensus subprofile is defined, which then in turn could be used to determine the biological similarity of new compounds to a respective cluster (Figure 1A). For subprofile identification, the following procedure was applied:

**Figure 1.**
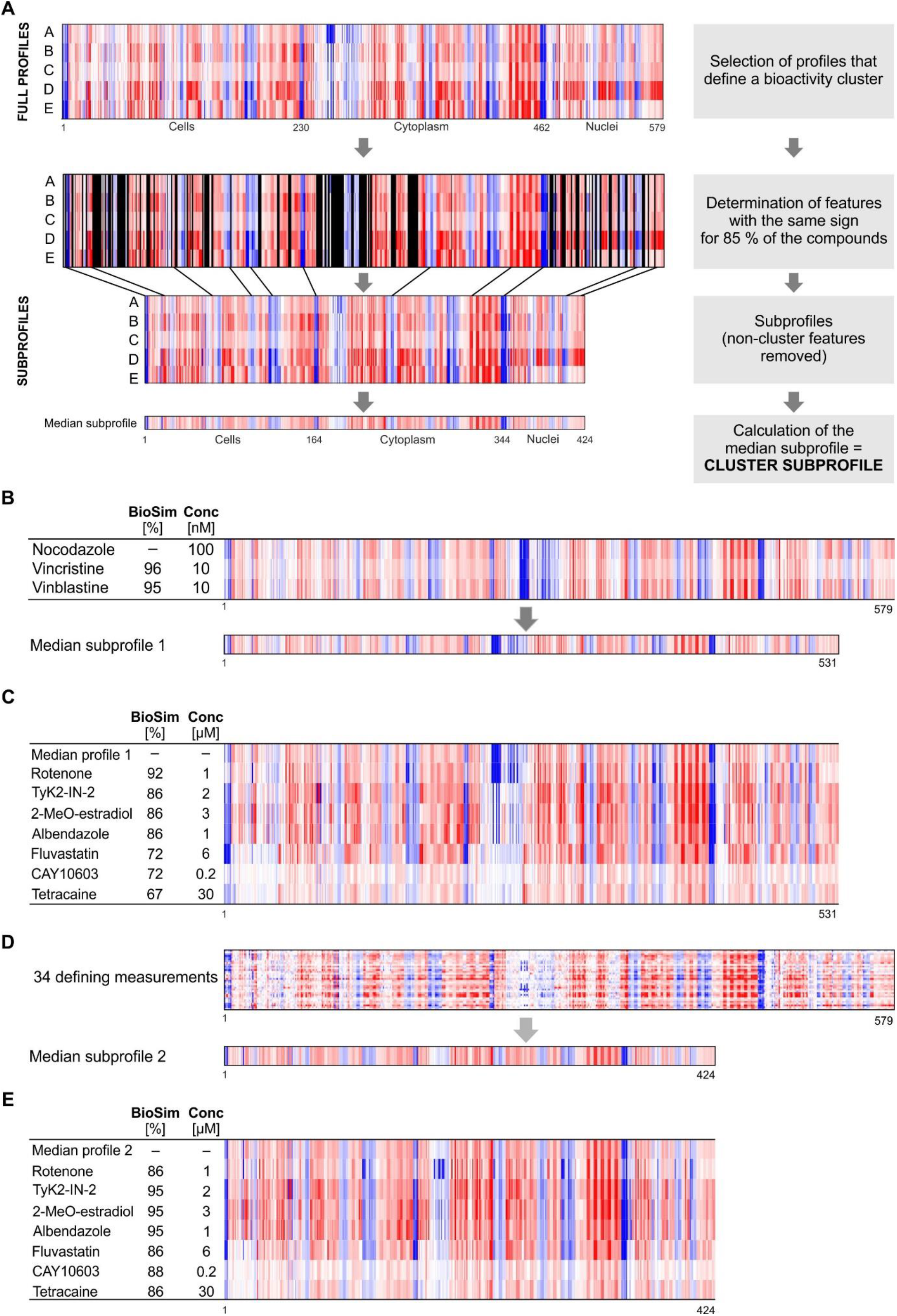
Extraction of cluster subprofiles. (A) Workflow for extracting relevant features from a set of defining features and calculation of median subprofile. From the full morphological profiles for biosimilar reference compounds (compounds A-E), features whose values do not have the same sign for 85% of the defining compounds will be removed (larger areas of these features that will be removed are marked in black). This results in reduced (shorter) profiles, i.e. subprofiles, containing only the homogeneous features (here 424). (B-E) Extraction of Tubulin cluster subprofile. (B) Narrow definition of the cluster using only Nocodazole, Vincristine and Vinblastine. The high biosimilarity of these three compounds results in a long median cluster subprofile with 531 features (92 % of the full profile). (C) Biosimilarity to the cluster of known tubulin-targeting compounds shows a low similarity for some compounds. (D) Broader definition of the tubulin cluster using 34 profiles from 26 confirmed tubulin-targeting agents results in a shorter median cluster subprofile with 424 features. (E) Biosimilarity of the same compounds as in (C) now shows values ≥ 85 % for all examples. Blue color: decreased feature, red color: increased feature.

In Pseudo-code:

~~~
function cluster_features(list_of_cluster_defining_compounds):
    list_of_cluster_features = empty_list()
    number_of_defining_compounds =
        length(list_of_cluster_defining_compounds)
    for each feature in all_features:
        count_plus = 0
        count_minus = 0
    for each compound in list_of_cluster_defining_compounds:
        if feature_value_of(compound) >= 0:
            count_plus += 1
        else:
            count_minus += 1
    fraction = max(count_plus, count_minus) /
        number_of_defining_compounds
    if fraction >= 0.85:
        list_of_cluster_features.append(feature)
    return list_of_cluster_features
~~~

(The Python implementation can be found in the function jupy_tools.cpa.cluster_features of the Github repository https://github.com/mpimp-comas/2022_pahl_ziegler_subprofiles)

From the list of cluster features subsequently a representative median subprofile for the cluster can be calculated by taking the median values over all compounds from the defining set for every given feature and combining them into a new reduced profile (Figure 1A). This median (*consensus*) subprofile can then be used for the calculation of similarity of test compounds to the cluster (Figure 1A). The dominating features for a cluster are then defined as all features of the full Cell Painting profile, where the values of 85% of the compounds from the cluster-defining set have the same sign (are all either positive or negative).

Initially we extracted the subprofile features for the tubulin cluster based on the three highly similar compounds Vinblastine, Vincristine and Nocodazole (Akbarzadeh et al., 2021). The resulting median subprofile consisted of 531 features instead of 579 features of the full profiles (Figure 1B). We then searched for reference compounds that are biosimilar to the tubulin cluster subprofile by calculating the similarity only between the extracted 531 features for all compounds. Several known tubulin-targeting small molecules displayed subprofile similarity of more than 85%, e.g., Rotenone, Tyk2-IN-2, 2-Methoxy-estradiol, and Albendazole (Figure 1C). However, the subprofiles for Fluvastatin (6 µM), CAY10603 (0.2 µM) and Tetracaine (30 µM), which are reported to impair microtubules (Yoon et al., 2011, Ma et al., 2019, Ali et al., 2013), showed similarity lower than 75% to the tubulin cluster subprofile (see Figure 1C). As the profiles for Vinblastine, Vincristine and Nocodazole are highly similar (≥ 95%), the tubulin cluster subprofile contains 92% of all features. Considering that many references that target tubulin have a nominal target different from tubulin (the nominal target is the target that is commonly associated with the compound (Moret et al., 2019)), their profiles may also contain the morphological signature that is caused by the nominal target, which may partly overlay the tubulin subprofile. To account for polypharmacology responses in the Cell painting analysis, we selected a diverse set of 34 profiles from 26 confirmed tubulin-targeting agents, many of which are known to target also different proteins (see cluster ‘Tubulin’ in Table S1), to generate the tubulin signature (Akbarzadeh et al., 2021). The resulting tubulin cluster subprofile includes 424 features, i.e., 107 features less than the initially created tubulin cluster profile based only on Vinblastine, Vincristine and Nocodazole (Figure 1D). Using this tubulin cluster subprofile, we detected biosimilarities to the tubulin cluster of higher than 85% for Fluvastatin, CAY10603 and Tetracaine, which would confirm the compounds as tubulin-targeting agents (Figure 1E).

We applied the same strategy to extract subprofiles of clusters that are based on modulation of AKT/PI3K/MTOR, Aurora kinase, BET, DNA synthesis, HDAC, HSP90, lysosomotropism/cholesterol homeostasis, protein synthesis or uncoupling of the mitochondrial proton gradient. For this purpose, for each of the clusters, a set of compounds with confirmed activity was defined (see Table S1 for the respective set of reference compounds that were used to define the clusters for subprofile extraction).

We then extracted the cluster features and generated the cluster subprofiles of the ten identified biological clusters, which were of different length (see Figure 2A): whereas the DNA synthesis or Aurora kinase subprofile are characterized by 288 and 358 features, respectively, the lysosomotropism/cholesterol homeostasis (L/CH) cluster subprofile contains 504 features. A biosimilarity search using the obtained subprofiles successfully identified the references of the respective compound set for each cluster (Table S1). These ten clusters were mapped in a lower dimension plot, using Uniform Manifold Approximation and Projection (UMAP (McInnes et al., 2018), see Figure 2B). Several clusters, although separated from each other, were located close together as also demonstrated by cross-correlation analysis (Figure 2C and Figure S2), which may complicate or even prevent MoA assignment for compounds based only on the location in the plot.

**Figure 2.**
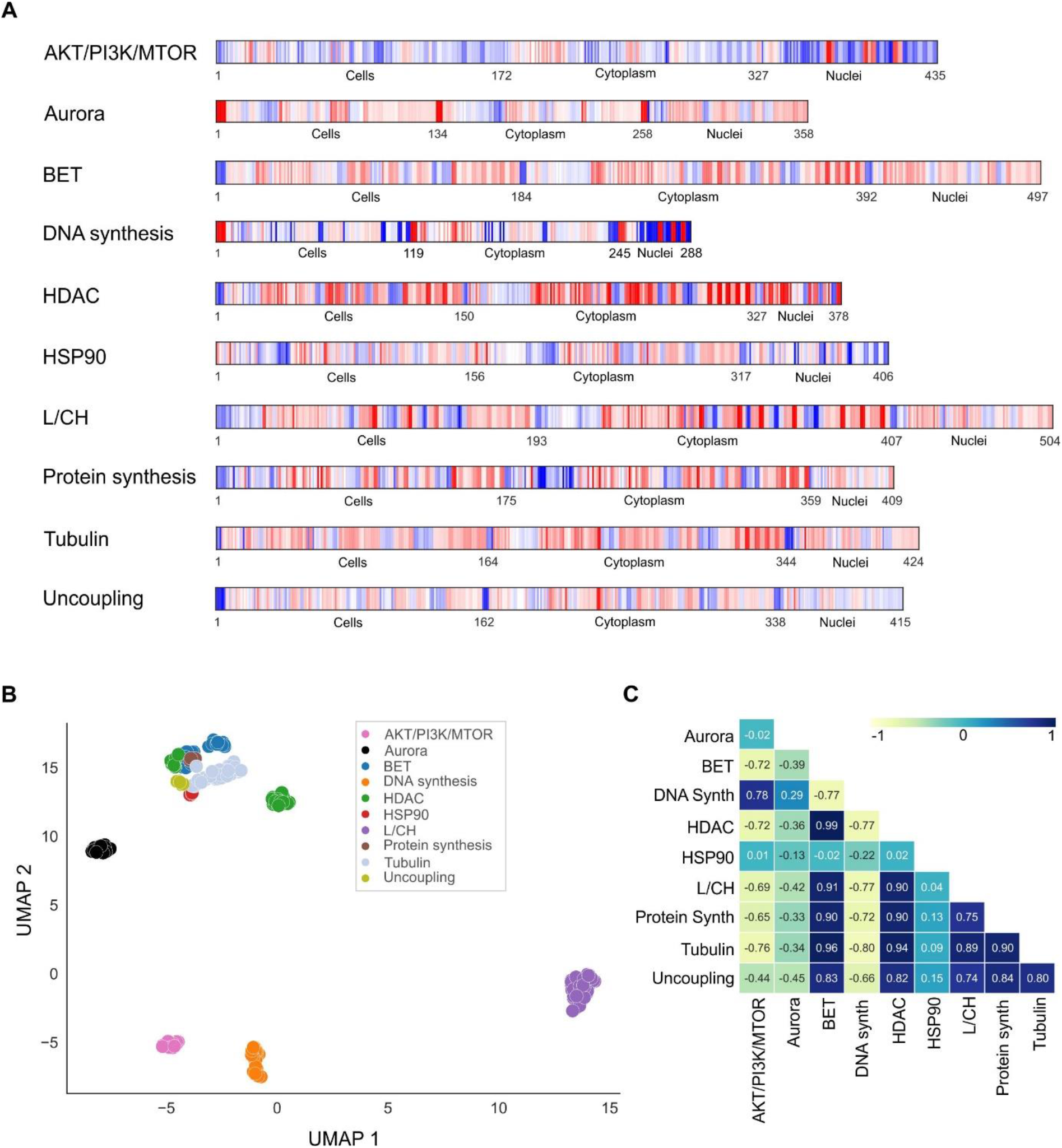
Subprofiles of the ten defined bioactivity clusters. (A) Median cluster subprofiles of the ten defined clusters. Blue color: decreased feature, red color: increased feature. (B) UMAP plot using the full profiles of the reference compounds that were used to define the bioactivity clusters. Not normalized data, 15 neighbors. (C) Cluster subprofile cross-correlation using Pearson correlation. L/CH: lysosomotropism/cholesterol homeostasis cluster.

Shared subprofile biosimilarity is expected to uncover the mode of action for unexplored compounds or to find unanticipated activity for reference compounds. For instance, a biosimilarity search using the tubulin cluster subprofile identified the PDGFRβ and B-Raf inhibitor KG 5 and the JAK inhibitor AZ 960 as potential microtubule-interfering compounds (Figure 3A). The cluster biosimilarity heatmap revealed high similarity to the tubulin cluster (Figure 3B). Whereas the full profile of KG 5 shares 94% biosimilarity to the profile of Nocodazole, the full profile similarity for AZ 960 is 75 %, and, thus, at the lower limit (Figure 3C and 3D). However, the subprofile analysis clearly indicates similarity of AZ 960 to the tubulin cluster (cluster similarity of 90%). Indeed, upon treatment with the compounds, microtubule organization was disturbed (AZ 960) or microtubules were depolymerized (KG 5) (Figure 3E). These results confirm impairment of the tubulin cytoskeleton by KG 5 and AZ 960 at low micromolar concentrations, which should be considered when these compounds are used to modulate their nominal targets. Subprofile analysis predicted HDAC-related activity for two yet unexplored compounds (compound **1** and **2**, Figure 4A and 4B) both bearing a trifluoromethyl ketone. This moiety is present in some HDAC inhibitors and is involved in zinc (II) coordination (Frey et al., 2002). Compound **1** and **2** differ by only one methylene group and share high biosimilarity with the HDAC inhibitor trichostatin A (TSA, see Figure 4C) and to the HDAC cluster subprofile (Figure 4B and 4D). Therefore, the influence of **1** and **2** on HDAC activity was explored in nuclear extracts from HeLa cells. Both compounds inhibited *in vitro* deacetylation by ca. 30 % (Figure 4E) at 30 µM. When U2OS cells were incubated with the compounds for 2 h, HDAC activity was reduced down to ca. 20 % (Figure 4F), thus confirming inhibition of HDAC activity by **1** and **2**.

**Figure 3.**
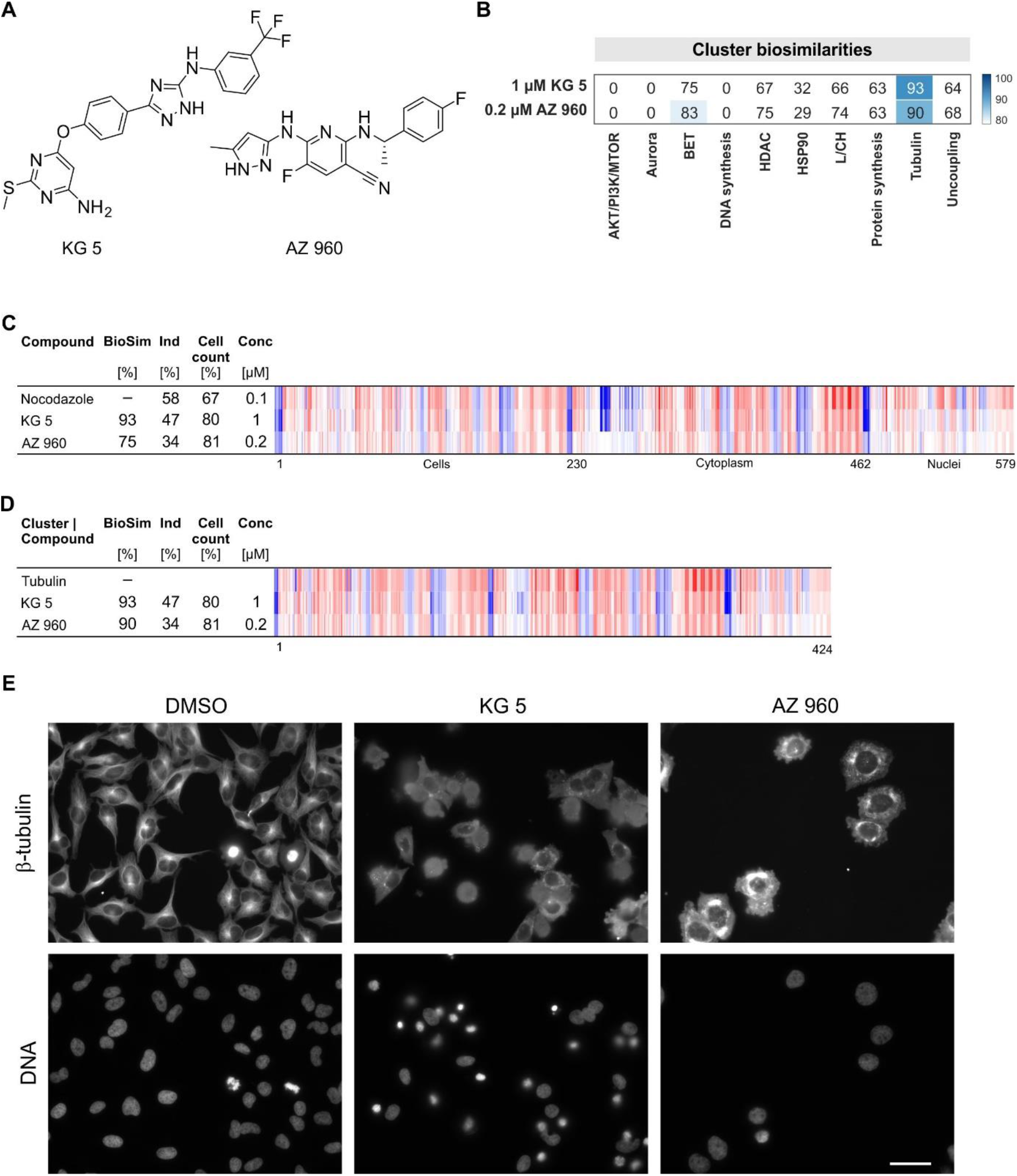
Identification of microtubule inhibitors. (A) Structures of KG 5 and AZ 960. (B) Cluster biosimilarity heatmap for KG 5 and AZ 960. (C) Biosimilarity of KG 5 and AZ 960 to Nocodazole. The top line profile is set as a reference profile (100 % biological similarity, BioSim) to which the following profiles are compared. Blue color: decreased feature, red color: increased feature. BioSim: biosimilarity, Ind: induction, Conc: concentration. (D) Biosimilarity of KG-5 and AZ 960 to the tubulin cluster subprofile. (E) Influence of KG 5 (10 µM) and AZ 960 (6 µM) on microtubules in U2OS cells. Cells were treated with the compounds for 24 h prior to staining for DNA and tubulin. Scale bar: 50 µm

**Figure 4:**
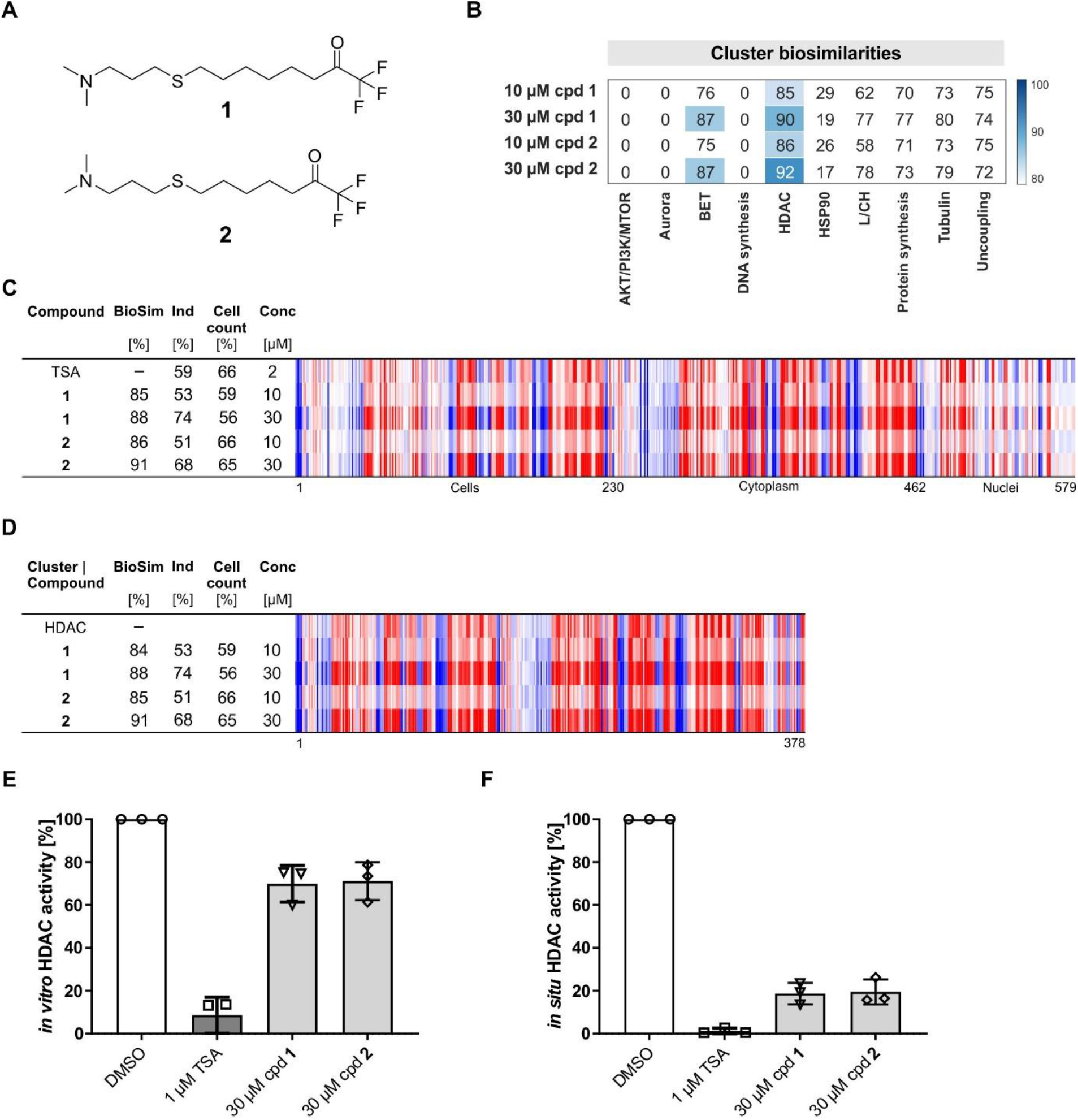
Inhibition of HDAC activity by compounds 1 and 2. (A) Structures of compounds **1** and **2**. (B) Cluster biosimilarity heatmap for compound **1** and **2**. (C) Biosimilarity of **1** and **2** to trichostatin A (TSA). The top line profile is set as a reference profile (100 % biological similarity, BioSim) to which the following profiles are compared. Blue color: decreased feature, red color: increased feature. BioSim: biosimilarity, Ind: induction, Conc: concentration. (D) Biosimilarity of **1** and **2** to HDAC cluster subprofile. (E) Influence on *in vitro* HDAC activity. (F) Influence on HDAC activity in HeLa cells upon treatment with the compounds for 24 h. Data are mean values (n = 3) ± SD.

Furthermore, the subprofile analysis identified the ALK2 and ALK3 inhibitor LDN193189 and macrocycle **3** as potential protein synthesis inhibitors (Figure 5A and 5B). At a concentration of 1 µM, LDN193189 shares 91 % subprofile similarity to the protein synthesis cluster (Figure 5B-5D). The subprofile of compound **3** (Dow et al., 2017) was 87 % similar to the subprofile of protein synthesis inhibitors (Figure 5B-5D). Indeed, both compounds inhibited *in vitro* protein translation (Figure S3A). Compound **3** and to a lesser extent LDN193189 suppressed protein synthesis also in cells (Figure 5E and 5F, higher concentrations than 1 µM of LDN193189 could not be used due to substantially reduced cell count). These results confirm inhibition of protein translation by LDN193189 and compound **3**. LDN193189 is widely used in stem cell culture to induce differentiation, albeit at lower concentrations than 1 µM (Konagaya and Iwata, 2019, Horbelt et al., 2015). To the best of our knowledge, no impairment of protein translation by LDN193189 has been reported thus far. However, gene expression analysis using L1000, a platform for exploring the expression of 1000 landmark genes that represent the transcriptome (Subramanian et al., 2017), suggests connectivity between LDN193189 and protein synthesis in HT29, MCF7 and PC3 cells (Figure S3B) that further supports our findings.

**Figure 5.**
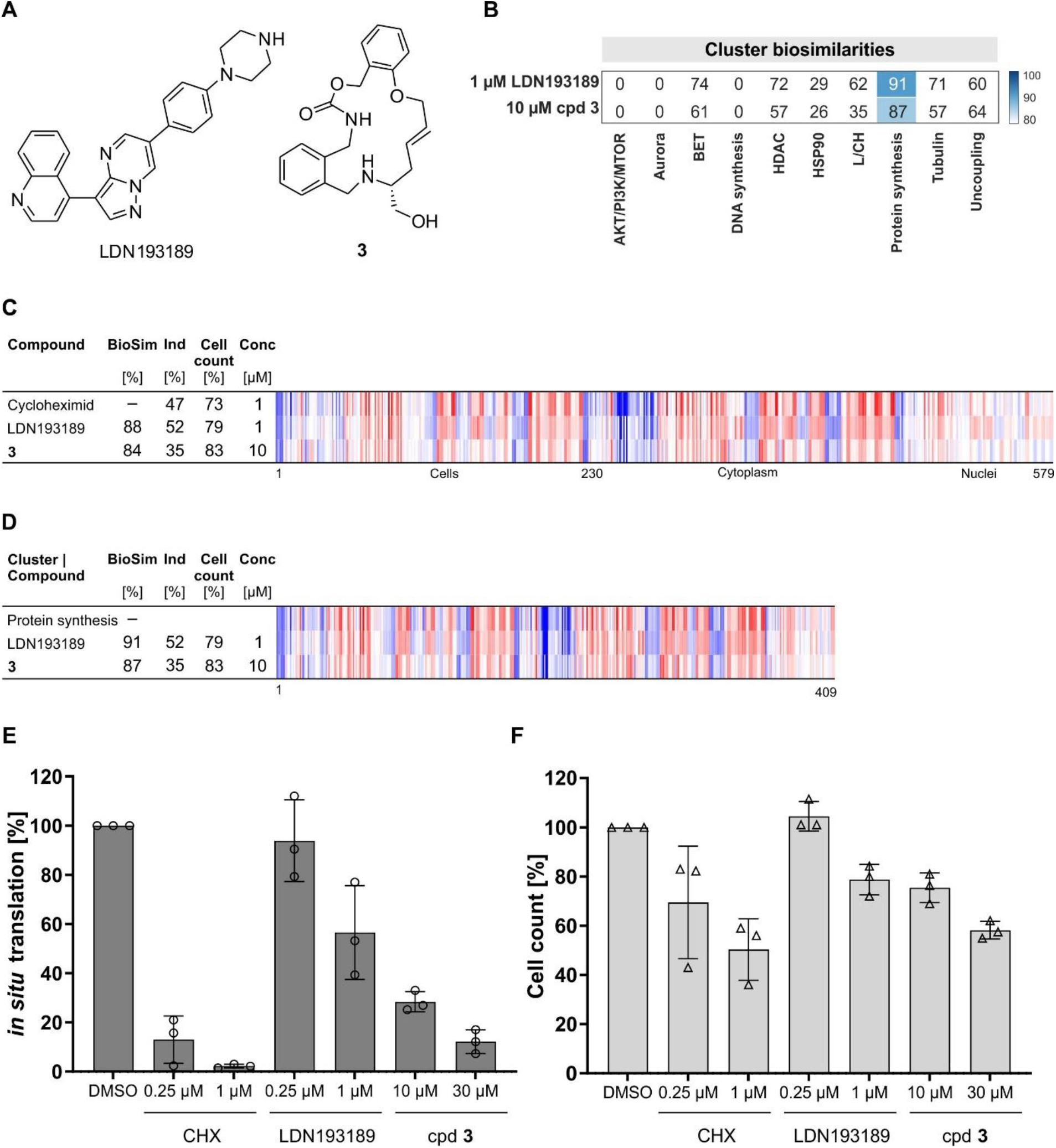
Identification of protein synthesis inhibitors. (A) Structures of the ALK2 and ALK3 inhibitor LDN193189 and compound **3**. (B) Cluster biosimilarity heatmap for LDN193189 and compound **3**. (C) Biosimilarity of LDN193189 and compound **3** to cycloheximid. The top line profile is set as a reference profile (100 % biological similarity, BioSim) to which the following profiles are compared. Values were normalized to the DMSO control. Blue color: decreased feature, red color: increased feature. The set of 579 features is divided in features related to the cell (1–229), cytoplasm (230–461) and nuclei (462–579). BioSim: biosimilarity, Ind: induction, Conc: concentration. (D) Biosimilarity of LDN193189 and compound **3** the protein synthesis cluster subprofile. (E and F) Influence on protein synthesis in HeLa cells. Cells were treated for 24 h with the compounds or cycloheximid (CHX) and DMSO as controls prior to detection of protein synthesis (D) or cell count (E). Data are mean values (n = 3) ± SD.

As a subprofile constitutes only part of the full profile, we used the non-cluster features to gain further bioactivity information. We previously assigned a large number of reference compounds which have very diverse annotated nominal targets, to the lysosomotropism/cholesterol homeostasis cluster (Schneidewind et al., 2021). Potentially, the nominal activity of these compounds may be detectable using CPA but masked by the dominant profile of disturbing cholesterol homeostasis. Therefore, we searched for biosimilar reference compounds using only the non-cluster features. For example, the NUAK1 inhibitor HTH-01-015 shares high similarity to the lysosomotropism/cholesterol homeostasis cluster at 10 µM but not at 3 µM (Figure 6A and 6B). However, bioactivity analysis for the HTH-01-015 at 10 µM using the subprofile of only the 75 non-cluster features revealed biosimilarity to the respective subprofiles of the Aurora kinase inhibitors SNS-314, ZM-447439 and Barasertib that are highly similar to the Aurora cluster, whereas the full profiles were dissimilar (Figure 6C and Figure S4A). Targeting of Aurora kinases by HTH-01-015 has not been demonstrated thus far (Banerjee et al., 2014). Analysis of Aurora kinase A, B and C activity using an *in vitro* kinase assay revealed dose-dependent inhibition of Aurora kinases with IC_50_ values of 5.6 µM (AURKA), 0.95 µM (AURKB) and 0.97 µM (AURKC), respectively. Of note, the cluster biosimilarity heatmap for HTH-01-015 at 3 µM suggests a potential inhibition of Aurora kinases (Figure 6B).

**Figure 6.**
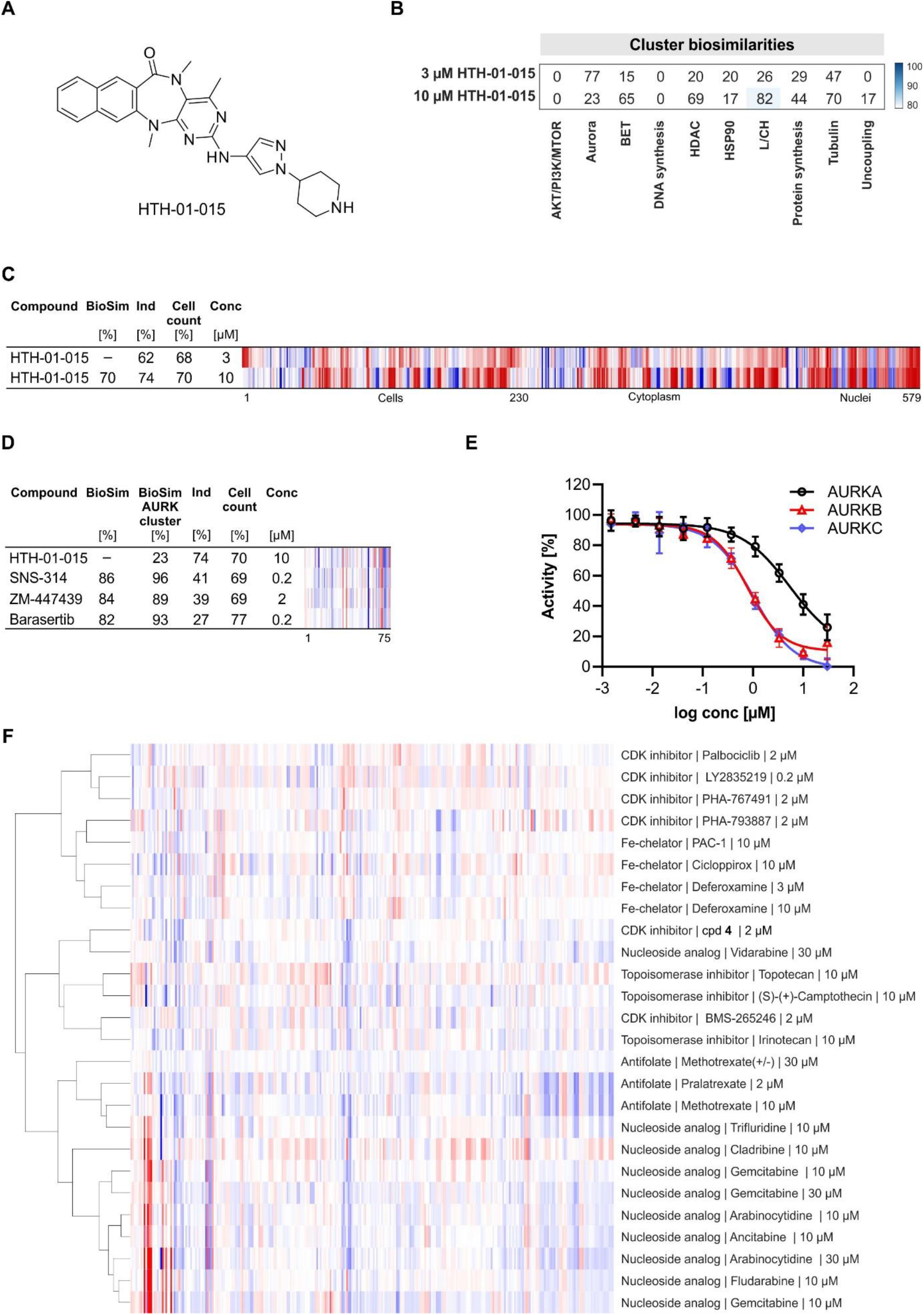
Non-cluster features for bioactivity assessment. (A) Structure of HTH-01-015. (B) Cluster biosimilarity heatmap for HTH-01-015. (C) Biosimilarity of HTH-01-015 profiles at 3 and 10 µM. (D) Biosimilarity of HTH-01-015 to Aurora inhibitors using only non-L/CH features. The top line profile is set as a reference profile (100 % biological similarity, BioSim) to which the following profile is compared. BioSim to Aurora cluster was calculated using the aurora cluster features. Blue color: decreased feature, red color: increased feature. BioSim: biosimilarity, Ind: induction, Conc: concentration. (E) Dose-dependent inhibition of Aurora kinases by HTH-01-015. Data are mean values (n = 2) ± SD. (F) Hierarchical clustering for subprofiles of compounds of the DNA synthesis cluster. Only non-cluster features were used. See Figure S5 for the structure of compound **4** (Bramson et al., 2001).

The full profiles for HTH-01-015 at 3 and 10 µM are not biosimilar (Figure 6C), whereas biosimilarity is detected when only the non-L/CH features are compared but not when L/CH or Aurora cluster subprofiles were used (Figure S4B-4D). HTH-01-015 has basic pKa of 10.08 and clogP of 2.28 and, thus, shares physicochemical properties with lysosomotropic compounds (lysosomotropic compounds typically have basic pKa > 6.5 and clogP > 2) (Nadanaciva et al., 2011). Lysosomotropic properties are often detected at higher concentrations, e.g. 10 µM, which explains the different morphological profiles for HTH-01-015 at 3 and 10 µM. These results demonstrate that non-cluster features can be employed for the detection of further bioactivities.

We envisioned that the non-profile features can further be used to differentiate between different mechanisms of action within a cluster that is based on a common mode of action, for instance the cluster of DNA synthesis/cell cycle inhibitors (Schneidewind et al., 2020). This cluster consists of compounds with different targets such as iron chelators, CDK and topoisomerase inhibitors as well as nucleosides and antifolates. Hierarchical clustering using the non-cluster profiles for compounds in this cluster revealed a clear separation of profiles of compounds sharing the same mechanism of action (see Figure 6F) and additionally suggest an unanticipated mechanism for compound **4** and BMS-265246, both reported to be CDK inhibitors.

## Discussion

Identification of small-molecule modulators of disease-relevant targets and processes is in focus of chemical biology research and drug discovery. Hit rates in typical target- and cell-based assays are often below 1 % and, thus, the majority of the screened compounds are inactive in the respective setup. Hence, there is a high demand for bioactivity annotations in compound screening collections, which additionally may facilitate target deconvolution in phenotypic screening. Conducting numerous target- and cell-based assays is time- and labor-consuming and still cannot cover the sheer variety of bioactivity that can be exerted by small molecules in cells. In contrast, profiling approaches, such as gene and protein expression profiling or morphological profiling, provide a more holistic view on the bioactivity of small molecules in cells by detecting hundreds of features. Morphological profiling can be performed in medium to high throughput and is thus particularly suitable for bioactivity detection. In contrast to transcriptomics and proteomics, linking altered morphological features to upstream regulation is challenging or even impossible without additional information and data. Therefore, morphological profiling relies on the profiles of reference compounds for the generation of target or mode-of-action hypothesis. However, we and others noticed that profiles of reference compounds with similar target annotation can differ as the activity may be governed not by the nominal target but rather an off-target (Akbarzadeh et al., 2021, Schneidewind et al., 2021, Cox et al., 2020, Woehrmann et al., 2013, Breinig et al., 2015). Moreover, reference compounds with diverse annotation may form a morphological cluster that is based on the same mode of action rather than on the same target (Schneidewind et al., 2020). Assignment of bioactivity by simply referring to the annotated MoA of the top most similar compounds may be in many cases misleading. Thus, bioactivity clusters have to be defined *a priori* to allow bioactivity prediction for uncharacterized compounds, which may also reveal unanticipated activity for reference compounds. Mapping the location of a compound profile in a lower dimension plot may provide insight into the MoA if the profile is located in a predefined cluster. However, due to polypharmacology, profiles may locate in the space between the clusters (Akbarzadeh et al., 2021), thus hampering MoA assignment. To facilitate profile analysis, we introduce the concept of subprofiles that contain only features describing the bioactivity of a predefined cluster. Subprofiles are extracted from the full profiles of a set of reference compounds and are specific for each cluster. Thus far, we defined ten morphological clusters based on targeting AKT/PI3K/MTOR, Aurora kinases, BET, DNA synthesis, HDAC, HSP90, lysosomotropism/cholesterol homeostasis, protein synthesis, tubulin and uncoupling of the mitochondrial proton gradient. These cluster subprofiles are of different length and compile the features that are characteristic for each individual cluster. The generated cluster subprofiles can be employed to simultaneously assess the similarity of small-molecule profiles to thus far ten clusters and, thereby to rapidly and reliably predict bioactivity related to one or more clusters. Whereas for full profiles we consider compounds as biosimilar if their profiles share > 75 % biosimilarity, this threshold lies at 80-85% as subprofiles contain less features. We demonstrate that the assignment of bioactivity based on the cluster subprofile analysis can subsequently be experimentally confirmed as exemplified for three uncharacterized compounds targeting HDAC or protein synthesis, which were synthesized in a different context than the detected bioactivity.

The subprofile approach substantially simplifies the analysis of the multidimensional profiles from the Cell Painting assay and allows scientists with different backgrounds (e.g., chemists, biologists and computational scientists) to easily obtain mode-of-action information without the need to inspect all biosimilar profiles and references. Moreover, the subprofile analysis bypasses the issue with incomplete bioactivity annotation and polypharmacology that in many cases can be misleading and generate wrong conclusions. In addition, subprofile analysis may be superior to cluster analysis using dimensionality reduction as polypharmacology may be reflected in the profiles and such profiles would not be assigned to one given cluster in a lower dimension plot as recently reported for CF53 (Akbarzadeh et al., 2021).

Not only the cluster features can be used for bioactivity analysis but also the non-cluster features may provide further information for bioactivity annotation as compounds have more than one target. Impairment of more than one target may lead to complex, mixed profiles. Deconvolution of the underlying MoA of mixed profiles is challenging and the subprofile approach promises to simplify their analysis. Subprofile similarity search with only the non-cluster features may reveal a second target or MoA beyond the one that determines the respective cluster bioactivity as demonstrated for the NUAK1 inhibitor HTH-01-015, which CPA assigned to the lysosomotropism/cholesterol homeostasis cluster. Analysis of the non-cluster features suggested inhibition of Aurora kinases that was subsequently confirmed. Moreover, non-cluster features can be employed to explore the mechanisms of action of a cluster that unites compounds with different targets such as the DNA synthesis cluster. This strategy allows for even more precise target prediction that may significantly shorten the target validation process.

There is a widespread interest in academia and pharmaceutical industry for thorough bioactivity annotation of compound collections. Detailed knowledge of the bioactivity of compounds may guide the hit triage process for selecting the most promising hits in screens (Vincent et al., 2020). In this regard, cluster subprofile analysis will early on point towards potential targets of a hit compound, which may spur or prevent its priorization for further studies. Cluster subprofile analysis will complement structure-activity relationship (SAR) studies on a given target or process and may provide insights for the unexpected behavior of derivatives in cellular assays. Moreover, this approach may indicate an off-target that may account for the activity observed in cell-based assays and will assist correct data interpretation, which is essential for proper hit selection. To support the chemical biology community, we disclose the cluster biosimilarity for our reference compound set that is accessible via https://github.com/mpimp-comas/2022_pahl_ziegler_subprofiles and the web app tool https://cpcse.pythonanywhere.com/.

In summary, we employed morphological subprofiles to define consensus cluster features and, thus, cluster subprofiles. Cluster subprofile analysis allows for easy and fast annotation of thus far ten different modes of action for compound collections without the need for exploring the profiles of the most similar reference compounds. The remaining non-cluster features may guide the mapping of further bioactivities and enable differentiation between mechanisms of action within a cluster that is based on a common mode of action.

## Methods

### Material

Dulbecco’s Modified Eagle’s medium (DMEM), L-glutamine, sodium pyruvate and non-essential amino acids were obtained from PAN Biotech, Germany. Roswell Park Memorial Institute (RPMI) 1640 medium without methionine and fetal bovine serum (FBS) was obtained from Gibco, Thermo Fisher Scientific Inc., USA. Anti alpha tubulin-FITC mouse mAb (#F2168) and 4’,6-diamidine-20-phenylindole dihydrochloride (DAPI, #10236276001) were purchased from Sigma Aldrich, Germany. Phospho-Histone H3 (Ser10) (D2C8) XP® Rabbit mAb Alexa Fluor® 594 Conjugate (#8481) was purchased from Cell Signaling Technology Europe, Germany. HDAC Activity Assay Kit (#566328) and *In Situ* HDAC Activity Fluorometric Assay Kit (#EPI003) were purchased from Sigma Aldrich, Germany. Click-iT™ HPG Alexa Fluor™ 488 Protein Synthesis Assay Kit (#C10428) and 1-Step Human Coupled IVT Kit – DNA (#88882) were obtained from Thermo Fisher Scientific Inc., USA. Microplates (384 well, white, low volume #3826, 96 well; black, clear bottom #3340) were obtained from Corning, Sigma Aldrich, Germany, and glass bottom plates (P96-1-N) from Cellvis, USA.

### Cell lines

U2OS cells (Cat#300364, Cell Line Service, Germany) and HeLa cells (Cat# ACC 57 DSMZ, Germany) were cultured in DMEM supplemented with 10% FBS, 2 mM L-glutamine, 1 mM sodium pyruvate and non-essential amino acids. Cells were maintained at 37 °C and 5% CO_2_ in a humidified atmosphere. Cell lines were regularly assayed for mycoplasma and were confirmed to be mycoplasma-free.

### Cell Painting assay

The described assay follows closely the method described by (Bray et al., 2016). Initially, 5 µl U2OS medium were added to each well of a 384-well plate (PerkinElmer CellCarrier-384 Ultra). Subsequently, U2OS cell were seeded with a density of 1600 cells per well in 20 µl medium. The plate was incubated for 10 min at the ambient temperature, followed by an additional 4 h incubation (37 °C, 5% CO2). Compound treatment was performed with the Echo 520 acoustic dispenser (Labcyte) at final concentrations of 10 µM, 3 µM or 1 µM. Incubation with compound was performed for 20 h (37 °C, 5% CO2). Subsequently, mitochondria were stained with Mito Tracker Deep Red (Thermo Fisher Scientific, Cat. No. M22426). The Mito Tracker Deep Red stock solution (1 mM) was diluted to a final concentration of 100 nM in prewarmed medium. The medium was removed from the plate leaving 10 µl residual volume and 25 µl of the Mito Tracker solution were added to each well. The plate was incubated for 30 min in darkness (37 °C, 5% CO2). To fix the cells 7 µl of 18.5 % formaldehyde in PBS were added, resulting in a final formaldehyde concentration of 3.7 %. Subsequently, the plate was incubated for another 20 min in darkness (RT) and washed three times with 70 µl of PBS. (Biotek Washer Elx405). Cells were permeabilized by addition of 25 µl 0.1% Triton X-100 to each well, followed by 15 min incubation (RT) in darkness. The cells were washed three times with PBS leaving a final volume of 10 µl. To each well 25 µl of a staining solution were added, which contains 1% BSA, 5 µl/ml Phalloidin (Alexa594 conjugate, Thermo Fisher Scientific, A12381), 25 µg/ml Concanavalin A (Alexa488 conjugate, Thermo Fisher Scientific, Cat. No. C11252), 5 µg/ml Hoechst 33342 (Sigma, Cat. No. B2261-25mg), 1.5 µg/ml WGA-Alexa594 conjugate (Thermo Fisher Scientific, Cat. No. W11262) and 1.5 µM SYTO 14 solution (Thermo Fisher Scientific, Cat. No. S7576). The plate is incubated for 30 min (RT) in darkness and washed three times with 70 µl PBS. After the final washing step, the PBS was not aspirated. The plates were sealed and centrifuged for 1 min at 500 rpm.

The plates were prepared in triplicates with shifted layouts to reduce plate effects and imaged using a Micro XL High-Content Screening System (Molecular Devices) in 5 channels (DAPI: Ex350-400/ Em410-480; FITC: Ex470-500/ Em510-540; Spectrum Gold: Ex520-545/ Em560-585; TxRed: Ex535-585/ Em600-650; Cy5: Ex605-650/ Em670-715) with 9 sites per well and 20x magnification (binning 2).

The generated images were processed with the *CellProfiler* package (https://cellprofiler.org/, version 3.0.0) on a computing cluster of the Max Planck Society to extract 1716 cell features per microscope site. The data was then further aggregated as medians per well (9 sites -> 1 well), then over the three replicates.

Further analysis was performed with custom *Python* (https://www.python.org/) scripts using the *Pandas* (https://pandas.pydata.org/) and *Dask* (https://dask.org/) data processing libraries as well as the *Scientific Python* (https://scipy.org/) package.

From the total set of 1716 features, a subset of highly reproducible and robust features was determined using the procedure described by (Woehrmann et al., 2013) in the following way: Two biological repeats of one plate containing reference compounds were analysed. For every feature, its full profile over each whole plate was calculated. If the profiles from the two repeats showed a similarity >= 0.8 (see below), the feature was added to the set.

This procedure was only performed once and resulted in a set of 579 robust features out of the total of 1716 that was used for all further analyses.

The phenotypic profiles were compiled from the Z-scores of all individual cellular features, where the Z-score is a measure of how far away a data point is from a median value.

Specifically, Z-scores of test compounds were calculated relative to the Median of DMSO controls. Thus, the Z-score of a test compound defines how many MADs (Median Absolute Deviations) the measured value is away from the Median of the controls as illustrated by the following formula:

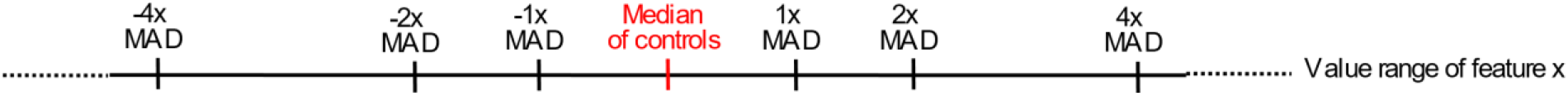

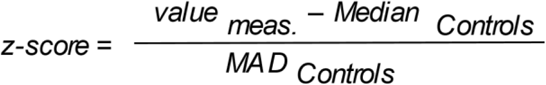

The phenotypic compound profile is then determined as the list of Z-scores of all features for one compound.

In addition to the phenotypic profile, an induction value was determined for each compound as the fraction of significantly changed features, in percent:

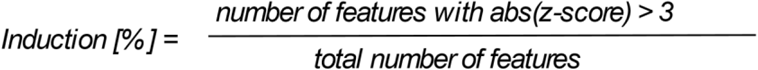

Similarities of phenotypic profiles were calculated from the correlation distances between two profiles(https://docs.scipy.org/doc/scipy/reference/generated/scipy.spatial.distance.correlation.html; Similarity = 1 - Correlation Distance), using either the feature values from the full profiles, those of the respective cluster subprofiles or those of the non-cluster profiles..

### Immunocytochemistry

5,000 U2OS cells were seeded per well in a black 96-well plate with clear bottom and incubated overnight. Cells were treated with compounds or DMSO as a control for 24 hours. Cells were then fixed using 3.7 % paraformaldehyde in phosphate-buffered saline (PBS) and permeabilized with 0.1% Triton X-100 (in PBS) prior to blocking nonspecific binding with 2% bovine serum umin (BSA in PBS). Staining with DAPI to visualize DNA and anti-tubulin-FITC antibody was performed in blocking buffer overnight at 4 C°. Images were acquired using Observer Z1 (Carl Zeiss, Germany) using 40x objective (LD Plan-Neofluar).

### *In vitro* HDAC Activity Assay

HDAC activity was assayed *in vitro* using the Histone deacetylase activity assay kit (#566328, Sigma Aldrich) according to the manufacturer’s protocol. 6-9 mg protein/ml HeLa Cell Nuclear Extract (Kit component No. KP31841 in 0.1 M KCl, 20 mM HEPES/NaOH pH 7.9, 20% (v/v) glycerol, 0.2 mM EDTA, 0.5 mM DTT, 0.5 mM PMSF) was diluted 1:30 in assay buffer (50 mM Tris/HCl, pH 8.0, 137 mM NaCl, 2.7 mM KCl, 1 mM MgCl_2_) and acts as a source for HDAC enzyme activity. The assay was performed in a white 384-well plate. A12.5µl reaction mixture contained 6.2 5µl substrate (Kit Component No. KP31842, 100 µM in assay buffer), which contains an acetylated lysine side chain, 3.75 µl HeLa Cell Nuclear Extract and 2.5 µl test compound or DMSO or Trichostatin A (1µM). The mixture was incubated for 30 min at room temperature. 12.5 µl developing solution and Trichostatin A (2 µM) were added to the samples for 10-20 min to convert the deacetylated substrate to a fluorophore. Fluorescence intensity was measured at ex/em 360/460 nm using a Tecan Spark plate reader.

### *In Situ* Histone Deacetylase (HDAC) Activity Fluorometric Assay

The assay was performed using the *In Situ* HDAC Activity Fluorometric Assay Kit (#EPI003, Sigma Aldrich) according to the manufacturer’s instructions. 20,000 U2OS cells per well were seeded into a black 96-well plate (clear bottom) and incubated overnight at 37°C and 5% CO_2_. The next day, cells were incubated for 2 h with the compounds or DMSO as a control together with a cell-permeable HDAC substrate, which contains an acetylated lysine side chain and a fluorophore that is quenched when bound to the substrate. Developer solution is added to lyse the cells and cleave the deacetylated HDAC substrate to release the fluorophore. After incubation for 30 min, fluorescence intensity was measured at ex/em 368/442 nm using the Tecan Spark plate reader.

### Click-it HPG Alexa Fluor Protein Synthesis Assay Kit

The assay was performed using the Click-iT™ HPG Alexa Fluor™ 488 Protein Synthesis Assay Kit (# C10428, Sigma Aldrich) according to the manufacturer’s instructions. 5,000 HeLa cells per well were seeded into a black 96-well plate (clear bottom) and incubated overnight at 37°C and 5% CO_2_.The next day, cells were treated with compound or DMSO as a control for 24 h. For degrading already synthesized protein cells were washed three times with phosphate-buffered saline (PBS) and incubated in methionine-free medium (RPMI 1640) supplemented with 10 % FBS for 30 min in the presence of compound or DMSO as a control. L-homopropargylglycine incorporation was started by adding Click-iT® HPG reagent (50 µM) in methionine-free medium (RPMI 1640) supplemented with 10 % FBS for 45 min. Fixation, permeabilization and Click-iT® HPG detection were performed according to the manufacturer’s protocol. HCS NuclearMask™ BlueStain was used to visualized DNA. Axiovert 200M microscope (Carl Zeiss, Germany) equipped with 10x objective was used to quantify Alexa Fluor® 488 using the software MetaMorph 7. Protein synthesis was assessed by determining signal intensity considering the total cell count as determined using the DNA.

### 1-Step Human Coupled IVT Kit

The assay was performed using the 1-Step Human Coupled IVT Kit (# 88882, Thermo Fisher Scientific Inc.) according to the manufacturer’s instructions using ‘turbo-type’ green fluorescent protein (tGFP) mRNA (# 88880 Thermo Fisher Scientific Inc.) as a template for translation. 5 µl HeLa lysate was preincubated with 1 µl of accessory protein for 10 min at room temperature in PCR plates (HSP9601, Bio-Rad). Reaction was started by adding 2 µl reaction mix, 1.2 µl tGFP mRNA (0.75 mg/µl) and 0.8 µl compound or DMSO as a control. Expression of tGFP was monitored for 5 h at 30°C using the CFX96 Touch Real-Time PCR Detection System (Bio-Rad Laboratories, Inc.).

### Quantification and statistical analysis

Data were either representative of three independent experiments or expressed as mean ± SD. All statistical details of the conducted experiments can be found in the respective figure caption. N: number of technical replicates, n: number of biological replicates.

### Data and code availability

The code to calculate subprofiles and profile biosimilarities, and to produce the plots in the figures can be found in the repository https://github.com/mpimp-\comas/2022_pahl_ziegler_subprofiles. The repository also contains the full processed Cell painting data set used in this analysis (3560 processed profiles at different concentrations). Cluster similarity for reference compounds can be accessed via the web app tool https://cpcse.pythonanywhere.com/.

## Acknowledgments

Research at the Max Planck Institute of Molecular Physiology was supported by the Max Planck Society. This work was co-funded by the European Union (Drug Discovery Hub Dortmund (DDHD), EFRE-0200481) and Innovative Medicines Initiative (grant agreement number 115489) resources of which are composed of financial contribution from the European Union’s Seventh Framework Programme (FP7/2007-2013) and EFPIA companies’ in-kind contribution, and by EPSRC (grants EP/N025652/1 and EP/F043503/1). Colin Ziegler is acknowledged for the design of the web app tool. Michael Grigalunas is acknowledged for the analysis of the chemistry data. The compound management and screening center (COMAS) in Dortmund is acknowledged for performing the high-throughput screening.

## Author Contributions

A.P. and S.Z. designed the research. B.S. performed the biological experiments. S.S. and A.P. performed and processed the CPA. M.R. and M.D. synthesized the compounds. A.N. and C.H. supervised the compound syntheses.. H.W. initiated and supervised the Cell painting study. A.P and S.Z. analyzed the CPA data. A.P. and S.Z. wrote the manuscript. All authors discussed the results and commented on the manuscript.

