## Supporting Information for "Morphological Subprofile Analysis for Bioactivity Annotation of Small Molecules"

### Supporting Figures

| Compound | BioSim [%] | Ind [%] | Cell count [%] | Conc [µM] |
|---|---|---|---|---|
| FCCP | – | 33 | 76 | 10 |
| AG 879 | 87 | 22 | 88 | 10 |
| A9 | 78 | 48 | 73 | 10 |
| Triclosan | 78 | 33 | 76 | 30 |
| IMD-0354 | 77 | 28 | 93 | 2 |

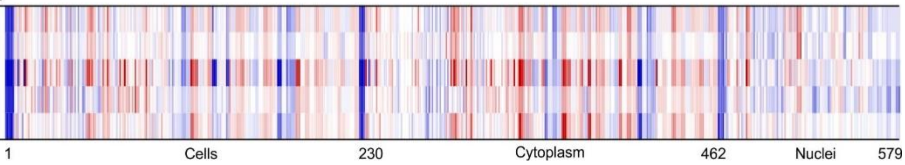


**Figure S1 (related to Figure 1). Profile similarity to FCCP.** The top line profile is set as a reference profile (100 % biological similarity, BioSim) to which the following profile are compared. Blue color: decreased feature, red color: increased feature. The set of 579 features is divided in features related to the cell (1–229), cytoplasm (230–461) and nuclei (462–579). BioSim: biosimilarity, Ind: induction, Conc: concentration. AG 879: Tyrphostin AG 879; A9: Tyrphostin A9.

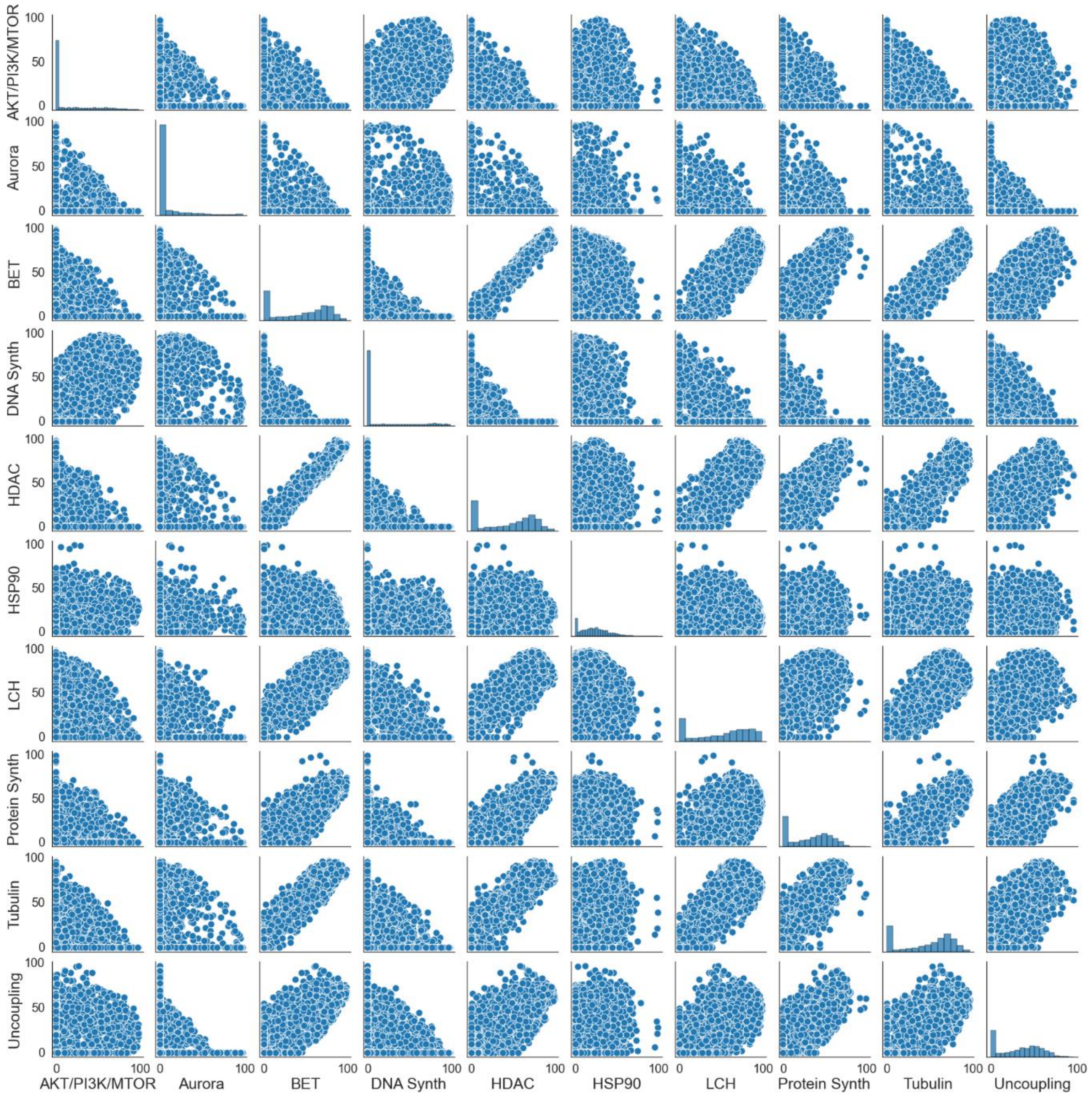


**Figure S2 (related to Figure 2). Pair plot of the cross-similarity analysis of the reference compounds to the ten defined clusters.** Each subplot contains the similarity to one biological cluster plotted against the similarity to another cluster for all 3,547 measurements of references considered in this study. The diagonal shows a histogram of the distribution for the respective similarity value.

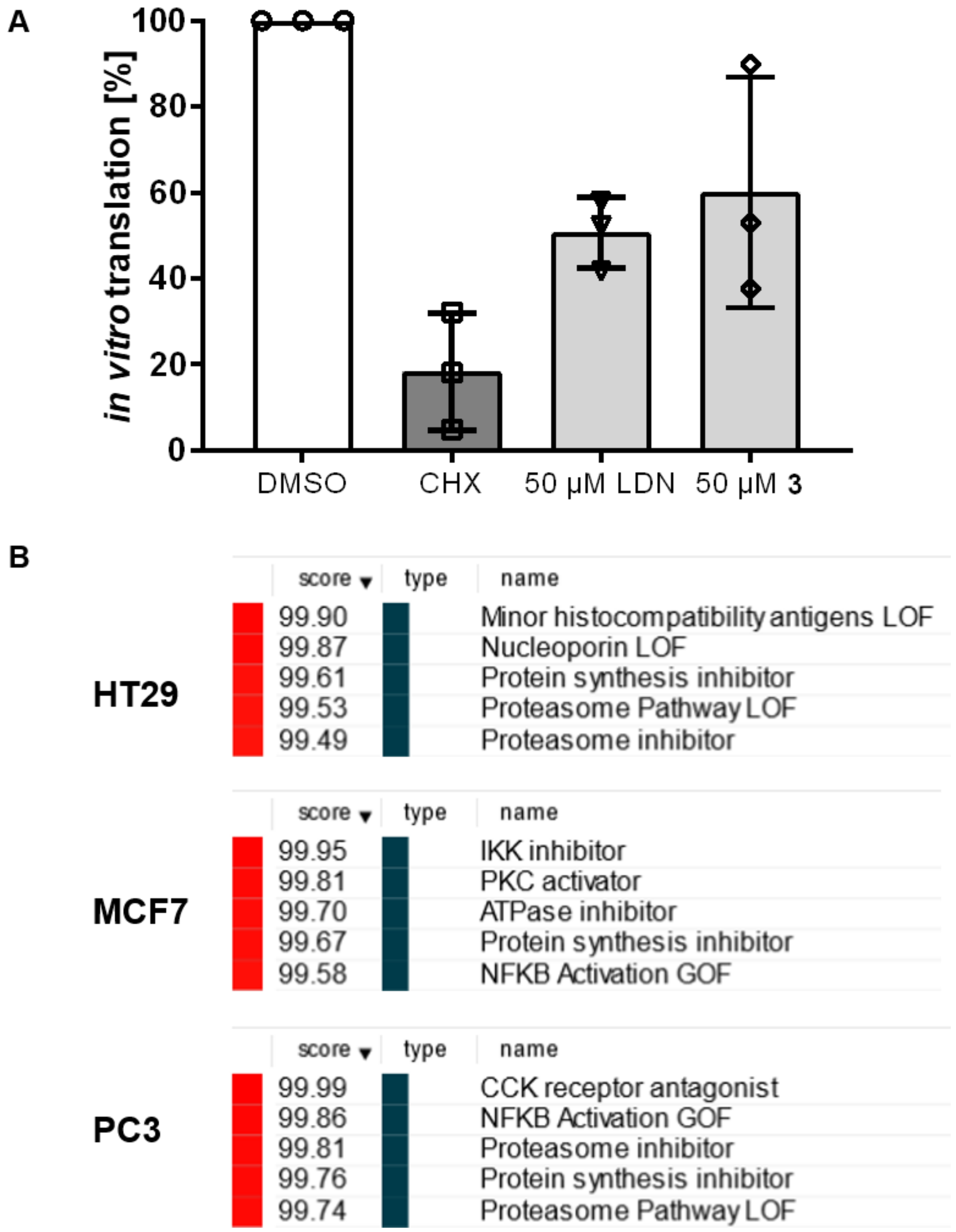


**Figure S3 (related to Figure 5). Influence on protein translation.** (A) Influence on of LDN193189 (LDN) and compound 3 protein translation *in vitro*. 10 µM cycloheximide (CHX) was used as a control. (B) Connectivty data for LDN193189 extracted using https://clue.io. Top five connectivities and the corresponding connectivity scores are displayed.

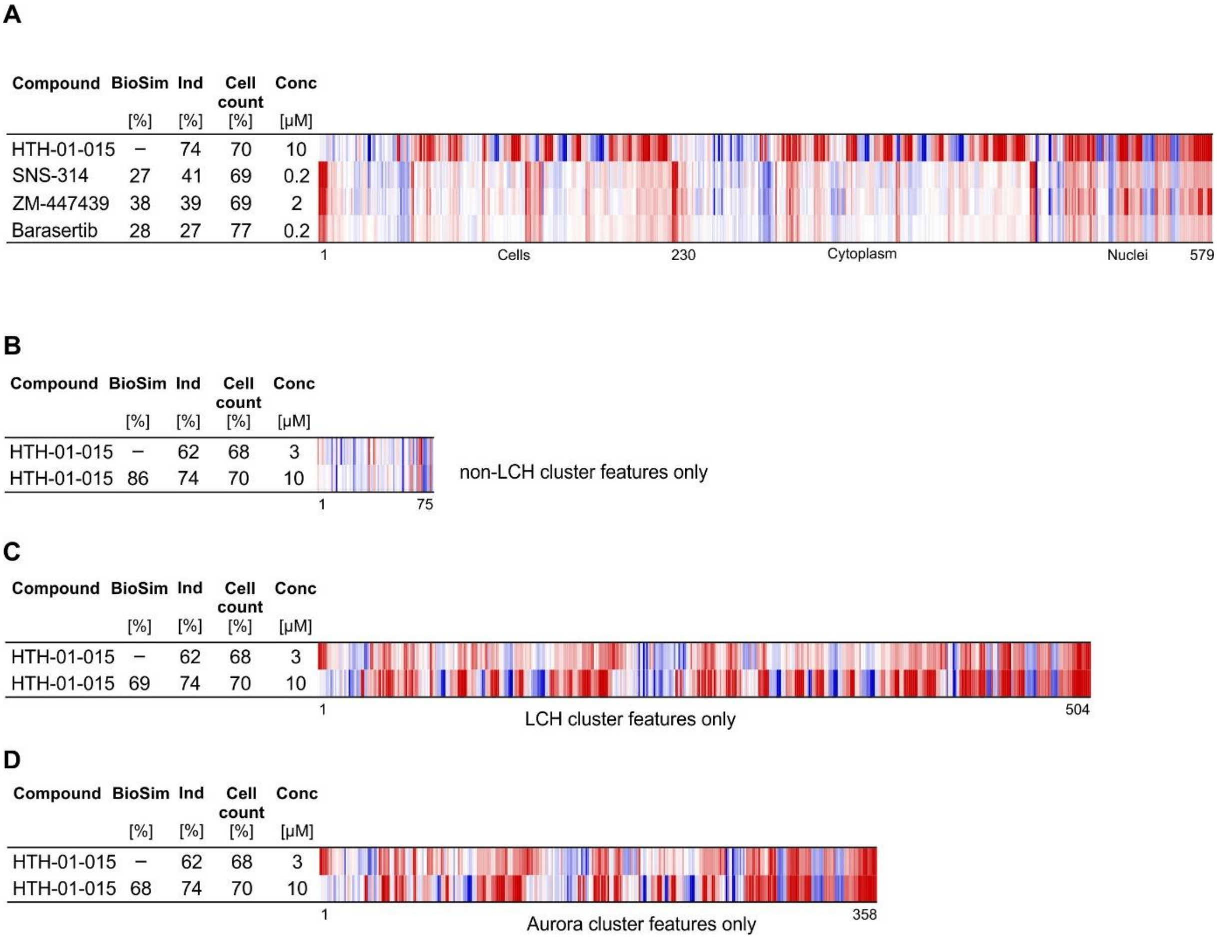


**Figure S4 (related to Figure 6). Profile analysis for HTH-01-015**. (A) Biosimilarity of HTH-01-015 to Aurora kinase inhibitors. (B-D) Biosimilarity of HTH-01-015 at 3 and 10 µM using only the non-L/CH cluster features (B), L/CH cluster features or (D) Aurora cluster features. The top line profile is set as a reference profile (100 % biological similarity, BioSim) to which the following profile is compared. BioSim to Aurora cluster was calculated using the aurora cluster features. Blue color: decreased feature, red color: increased feature. BioSim: biosimilarity, Ind: induction, Conc: concentration. (

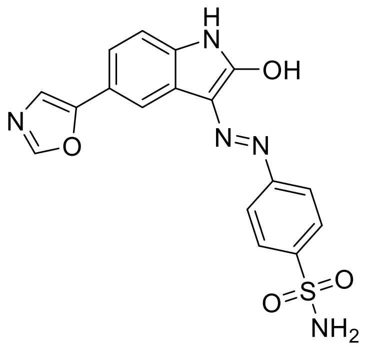


**Figure S5 (related to Figure 6). Structure of compound 4.**

### Chemistry

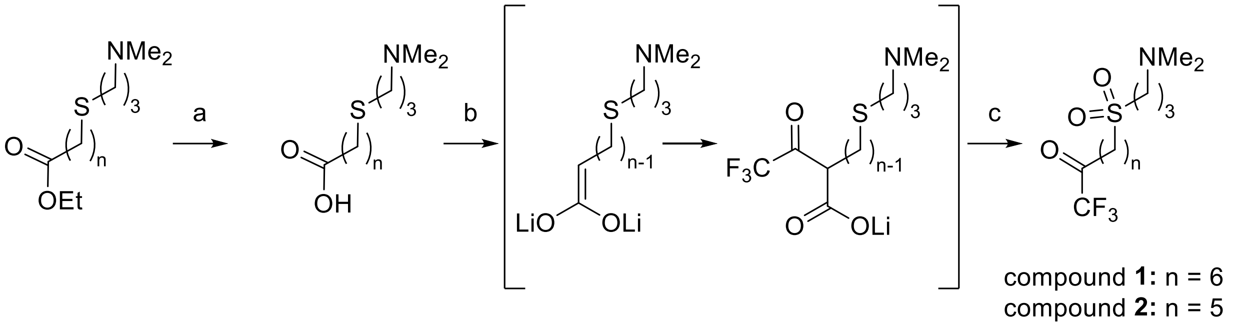


Synthesis of compounds **1** and **2**. Reagents and conditions: a) NaOH (1.0 eqv.), $H_2O$:EtOH, rt, 2h. b) Diisopropylamine (3.5 eqv.), n-Buli (3.4 eqv.), -78°C, carboxylic acid (1.0 eqv.), THF, rt (4h), $CF_3COOEt$ (3.0 eqv.,), -78°C (15 min), 6N HCl, 45-53 % yield (over two steps). c) Oxone® (3.0 eqv.), MeOH/$H_2O$ (3/2), overnight, rt, 24h, 35% yield.

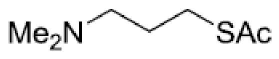


***N,N*-(Dimethylamino)propyl thioacetate** (**S1**)

Compound **S1** was synthesized according to Hedberg et al. (Hedberg et al., 2011)

**$^1$H NMR** (400 MHz, $CDCl_3$): δ 2.90 (t, $J$ = 7.2, 2H, -C$H_2$S), 2.31 (t, $J$ = 7.2, 2H, -C$H_2$N), 2.32 (s, 3H, COC$H_3$), 2.21 (s, 6H, N(C$H_3$)2), 1.74 (p, $J$ = 7.2, 2H). **$^{13}$C NMR** (125 MHz, $CDCl_3$): δ 195.7, 58.5, 45.5, 30.7, 27.8, 27.1. HRMS (ESI) calc. for $C_7H_{15}NOS$ [M+H]$^+$ 162.0947, found 162.0946. GCMS found 161 for [M$^+$.].

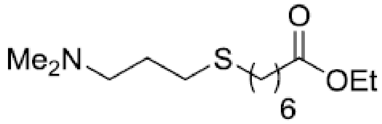

**Ethyl 7-(3-(dimethylamino)propylthio)heptanoate** (**S2**).

To a two-neck round bottom flask containing a stir bar and a solution of thioacetate **S1** (52 mmol, 1.0 eqv.) in anhydrous ethanol (80 ml) was added $Cs_2CO_3$ (58 mmol, 1.1 eqv.). The flask was equipped with a reflux condenser, and the reaction was flushed with Ar and kept under a positive pressure of Ar for the remainder of the reaction. The reaction was stirred at reflux until TLC analysis indicated that all of thioacetate **S1** had been consumed with the formation of the more polar thiol intermediate (EtOAc/CyHex 1/2). After this time, the reaction was cooled to 0 °C and a solution of ethyl 7-iodoheptanoate (58 mmol, 1.1 eqv.) in anhydrous ethanol (20 ml) was added dropwise. The reaction was then stirred at 22 °C for 1 h and then heated to 40 °C until the thiol intermediate was consumed (about 12 hours, TLC conditions: EtOAc/CyHex 1/2). After this time, the reaction was concentrated and EtOAc/$H_2O$ 1/2 (400ml) was added to the crude residue and stirred until a solution was formed. The phases were separated and the aq. layer was washed twice with EtOAc (100 ml). The organic layers were combined, washed with water (150 ml), brine (270 ml), dried over $MgSO_4$, and concentrated to afford compound **S2** without any further purification.

Yield = 75% (slight yellow oil). R*f* = 0.56 (AcOH/EtOAc/MeOH/$H_2O$ 3/3/3/2). IR: 1734 cm-1 (ester, C=O stretch). **$^1$H NMR** (400 MHz, $CDCl_3$): δ 4.10 (q, *J* = 7.2, 2H, -COC$H_2$CH$_3$), 2.52 (t, *J* = 7.2, 2H, -C$H_2$S),2.49 (t, *J* = 7.4, 2H, -C$H_2$S), 2.39 (t, *J* = 7.6, 2H, -C$H_2$COOEt), 2.27 (t, *J* = 7.6, 2H, -C$H_2$NMe$_2$), 2.25 (s,6H, N(C$H_3$)$_2$), 1.76 – 1.66 (m, 2H), 1.65 – 1.50 (m, 4H), 1.41–1.20 (m, 4H), 1.22 (t, *J* = 7.2, 3H, -COCH$_2$C$H_3$). **$^{13}$C NMR** (125 MHz, $CDCl_3$): δ 173.6, 60.1, 58.7, 45.4, 34.2, 32.1, 30.0, 29.4, 28.7, 28.5, 27.7, 24.8, 14.2. HRMS (ESI) calc. for $C_{14}H_{29}NO_2S$ $[M+H]^+$ 276.1992, found 276.1992. GCMS found 275 for $[M^{+}.]$.

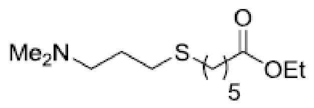

**Ethyl 6-(3-(dimethylamino)propylthio)hexanoate** (**S3**).

To a two-neck round bottom flask containing a stir bar and a solution of thioacetate **S1** (72 mmol, 1.0 eqv.) in anhydrous ethanol (108 ml) was added $Cs_2CO_3$ (78 mmol, 1.1 eqv.). The flask was equipped with a reflux condenser, and the reaction was flushed with Ar and kept under a positive pressure of Ar for the remainder of the reaction. The reaction was stirred at reflux until TLC analysis indicated that all of thioacetate **S1** had been consumed with the formation of the more polar thiol intermediate (EtOAc/CyHex 1/2). After this time, the reaction was cooled to 0 °C and a solution of ethyl 7-iodohexanoate (78 mmol, 1.1 eqv.) in anhydrous ethanol (27 ml) was added dropwise. The reaction was then stirred at 22 °C for 1 h and then heated to 40 °C until the thiol intermediate was consumed (about 12 hours, TLC conditions: EtOAc/CyHex 1/2). After this time, the reaction was concentrated and EtOAc/$H_2O$ 1/2 (540 ml) was added to the crude residue and stirred until a solution was formed. The phases were separated and the aq. layer was washed twice with EtOAc (150 ml). The organic layers were combined, washed with water (150 ml), brine (270 ml), dried over $MgSO_4$, and concentrated to afford compound **S3** without any further purification.

Yield = 77% (slight yellow oil). R*f* = 0.56 (AcOH/EtOAc/MeOH/$H_2O$ 3/3/3/2). IR: 1733 cm 1 (ester, C=O stretch). **$^1$H NMR** (400 MHz, $CDCl_3$): δ 4.01 (q, *J* = 7.2, 2H, -COC$H_2$CH$_3$), 2.40 (t, *J* = 7.4, 2H, -C$H_2$S), 2.39 (t, *J* = 7.4, 2H, -C$H_2$S), 2.21 (t, *J* = 7.2, 2H, -C$H_2$COOEt), 2.17 (t, *J* = 7.4, 2H, -C$H_2$NMe$_2$), 2.09 (s,6H, N(C$H_3$)$_2$), 1.65 – 1.57 (m, 2H), 1.55 – 1.43 (m, 4H), 1.34 – 1.24 (m, 2H), 1.13 (t, *J* = 7.2, 3H, -COCH$_2$C$H_3$). **$^{13}$C NMR** (125 MHz, $CDCl_3$): δ 173.2, 59.9, 58.4, 45.2, 33.9, 31.7, 29.7, 29.0, 28.1, 27.5, 24.3, 14.0. HRMS (ESI) calc. for $C_{13}H_{27}NO_2S$ $[M+H]^+$ 262.1835, found 262.1837. GCMS found 261 for $[M^+.]$.

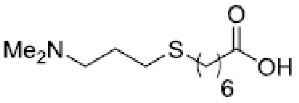


**7-(3-(Dimethylamino)propylthio)heptanoic acid** (**S4**).

To a 100 ml round bottomed flask equipped with a stir bar and a solution of ethyl ester **S2** (5.6 mml, 1 eqv. in 7 ml EtOH) was added 2 M NaOH (5.6 mmol, 1 eqv.). The reaction was stirred at 22 °C for two hours. Upon completion as determined by TLC (MeOH/DCM 2/8), the reaction was extracted with $Et_2O$ and the organic layer was discarded. The aq. layer was then acidified using conc. HCl to approximately pH 3. The aq. phase was then lyophilized, resulting in compound **S4** without the need for further purification. The product contains some NaCl salts.

Yellow solid. R*f* = 0.71 (AcOH/EtOAc/MeOH/$H_2O$ 3/3/3/2). IR: 1722 cm-1 (C=O). **1H NMR** (400 MHz, $CDCl_3$): δ 3.22 – 3.16 (m, 2H, -C*H*2COOH), 2.88 (s, 6H, N(C*H*3)2), 2.62 (t, *J* = 7.0, 2H, -C*H*2S), 2.56 (t, *J* = 7.4, 2H, -C*H*2S), 2.37 (t, *J* = 6.8, 2H, -C*H*2NMe2), 2.25 – 2.23 (m, 2H), 1.71 - 1.59 (m, 4H), 1.51 -1.36 (m, 4H). **$^{13}$C NMR** (125 MHz, $CDCl_3$): δ 176.9, 57.2, 43.4 (2C), 33.7, 30.8, 28.6, 28.2, 27.5, 27.2, 24.1, 23.9. HRMS (ESI) calc. for $C_{12}H_{25}NO_2S$ $[M+H]^+$ 248.1678, found 248.1679. LC-MS (ESI) found 248.04 for $[M+H]^+$.

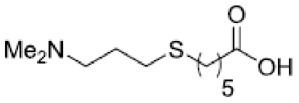


**6-(3-(Dimethylamino)propylthio)hexanoic acid** (S5).

To a 100 ml round bottomed flask equipped with a stir bar and a solution of ethyl ester **S3** (5.6 mml, 1 eqv. in 7 ml EtOH) was added 2 M NaOH (5.6 mmol, 1 eqv.). The reaction was stirred at 22 °C for two hours. Upon completion as determined by TLC (MeOH/DCM 2/8), the reaction was extracted with $Et_2O$ and the organic layer was discarded. The aq. layer was then acidified using conc. HCl to approximately pH 3. The aq. phase was then lyophilized, resulting in compound **S5** without the need for further purification. The product contains some NaCl salts.

Yellow solid. R*f* =0.61 (AcOH/EtOAc/MeOH/$H_2O$ 3/3/3/2). IR: 1729 cm-1 (C=O). **$^1$H NMR** (400 MHz, $CDCl_3$): δ 3.19 –3.13 (m, 2H, -C$H_2$COOH), 2.84 (s, 6H, N(C$H_3$)$_2$), 2.61 (t, *J* = 6.8, 2H, -C$H_2$S), 2.55 (t, *J* = 7.0, 2H, -C$H_2$S), 2.36 (t, *J* = 7.0, 2H, -C$H_2$NMe$_2$), 2.19 – 2.10 (m, 2H), 1.73 – 1.57 (m, 4H), 1.52 – 1.44 (m, 2H). **$^{13}$C NMR** (125 MHz, $CDCl_3$): δ 176.9, 57.1, 43.2 (2C), 33.7, 31.8, 28.7, 28.7, 27.7, 24.2, 24.0. HRMS (ESI) calc. for $C_{11}H_{23}NO_2S$ $[M+H]^+$ 234.1522, found 234.1522. LC-MS (ESI) found 234.05 for $[M+H]^+$.

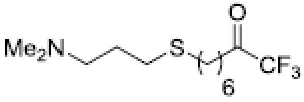


**8-(3-(Dimethylamino)propylthio)-1,1,1-trifluorooctan-2-one** (**1**).

Compound **S4** (0.63 mmol, 1.0 eqv.) was suspended in anhydrous THF (1.5 ml) and cooled to -20 °C. In a separate reaction flask, a solution of LDA was prepared by adding n-BuLi (2.5 M in hexane, 2.2 mmol, 3.4 eqv.) to a solution of freshly distilled DIPA (2.26 mmol, 3.5 eqv. in THF (1.2 ml)) at -78 °C. The solution of LDA was added dropwise over a period of 10 min to the solution of **S4** and was warmed to 22 °C and reacted for 4 hours. In a third flask, the enediolate solution was added dropwise to a solution of $CF_3CO_2Et$ (1.94 mmol, 3.0 eqv. in THF (0.7 ml)) at -78 °C. After stirring the reaction at -78 °C for 15 min, the reaction was quenched by adding 6M HCl (1.2 ml) dropwise. The reaction was diluted with EtOAc and the organic layer was isolated, dried ($MgSO_4$), and concentrated. The residue was purified by flash chromatography (1-8% MeOH in DCM) to afford compound **1**. Yield = 45% (over 2 steps, starting from ester **S2**, as yellow oil).

$R_f$ = 0.73 (AcOH/EtOAc/MeOH/$H_2O$ 3/3/3/2). IR: 1762 $cm^{-1}$ (C=O). **$^1$H NMR** (400 MHz, $CDCl_3$): δ 2.70 (t, *J* = 7.2, 2H, -C$H_2$COC$F_3$), 2.53 (t, *J* = 7.4, 2H, -C$H_2$S), 2.51 (t, *J* = 7.4, 2H, -C$H_2$S), 2.35 (t, *J* = 7.4, 2H, - C$H_2$N$Me_2$), 2.23 (s, 6H, N(C$H_3$)$_2$), 1.79 – 1.53 (m, 6H), 1.46 - 1.26 (m, 4H). **$^{13}$C NMR** (125 MHz, $CDCl_3$): δ 191.4 (q, CO, $J_2$ (C-F) = 34.6Hz), 115.5 (q, $CF_3$, $J_1$ (C-F)= 290 Hz), 58.5, 45.2 (2C), 36.2, 31.9, 29.9, 29.2, 28.3, 28.2, 27.3, 22.1. **$^{19}$F NMR** (377 MHz, $CDCl_3$): δ - 79.7 (s, $CF_3$). HRMS (ESI) calc. for $C_{13}H_{24}F_3NOS$ [M+H]$^+$ 300.1603, found 300.1606. GCMS found 299 for [$M^{+.}$].

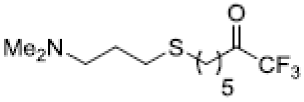


**7-(3-(Dimethylamino)propylthio)-1,1,1-trifluoroheptan-2-one** (**2**).

Compound **S5** (0.63 mmol, 1.0 eqv.) was suspended in anhydrous THF (1.5 ml) and cooled to -20 °C. In a separate reaction flask, a solution of LDA was prepared by adding n-BuLi (2.5 M in hexane, 2.2 mmol, 3.4 eqv.) to a solution of freshly distilled DIPA (2.26 mmol, 3.5 eqv. in THF (1.2 ml)) at -78 °C. The solution of LDA was added dropwise over a period of 10 min to the solution of **S4** and was warmed to 22 °C and reacted for 4 hours. In a third flask, the enediolate solution was added dropwise to a solution of $CF_3CO_2Et$ (1.94 mmol, 3.0 eqv. in THF (0.7 ml)) at -78 °C. After stirring the reaction at -78 °C for 15 min, the reaction was quenched by adding 6M HCl (1.2 ml) dropwise. The reaction was diluted with EtOAc and the organic layer was isolated, dried ($MgSO_4$), and concentrated. The residue was purified by flash chromatography (1-8% MeOH in DCM) to afford compound **2**. Yield = 52% (over 2 steps, starting from ester **S3**, as yellow oil).

$R_f$ = 0.67 (AcOH/EtOAc/MeOH/$H_2O$ 3/3/3/2). IR: 1762 $cm^{-1}$ (C=O). **$^1$H NMR** (400 MHz, $CDCl_3$): δ 2.72 (t, $J$ = 7.4, 2H, -C$H_2$COC$F_3$), 2.54 (t, $J$ = 7.2, 2H, -C$H_2$S), 2.52 (t, $J$ = 7.2, 2H, -C$H_2$S), 2.37 (t, $J$ = 7.2, 2H, - C$H_2$NMe$_2$), 2.24 (s, 6H, N(C$H_3$)$_2$), 1.81 – 1.57 (m, 6H), 1.49 - 1.39 (m, 2H). **$^{13}$C NMR** (125 MHz, $CDCl_3$): δ 191.3 (q, CO, $J_2$ (C-F) = 34.7 Hz), 115.48 (q, $CF_3$, $J_1$ (C-F)= 290 Hz), 58.5, 45.2 (2C), 36.2, 31.7, 29.9, 29.1, 27.8, 27.4, 21.9. **$^{19}$F NMR** (377 MHz, $CDCl_3$): δ - 79.7 (s, $CF_3$). HRMS (ESI) calc. For $C_{12}H_{22}F_3NOS$ [M+H]$^+$ 286.1447, found 286.1449. GCMS found 285 for [$M^{+\cdot}$].
